# Microbiome interaction networks and community structure from lab-reared and field-collected *Aedes aegypti, Aedes albopictus,* and *Culex quinquefasciatus* mosquito vectors

**DOI:** 10.1101/337311

**Authors:** Shivanand Hegde, Kamil Khanipov, Levent Albayrak, George Golovko, Maria Pimenova, Miguel A. Saldaña, Mark M Rojas, Emily A. Hornett, Greg C. Motl, Chris L. Fredregill, James A. Dennett, Mustapha Debboun, Yuriy Fofanov, Grant L. Hughes

## Abstract

Microbial interactions are an underappreciated force in shaping insect microbiome communities. Although pairwise patterns of symbiont interactions have been identified, we have a poor understanding regarding the scale and the nature of co-occurrence and co-exclusion interactions within the microbiome. To characterize these patterns in mosquitoes, we sequenced the bacterial microbiome of *Aedes aegypti, Ae. albopictus,* and *Culex quinquefasciatus* caught in the field or reared in the laboratory and used these data to generate interaction networks. For collections, we used traps that attracted host-seeking or ovipositing female mosquitoes to determine how physiological state affects the microbiome under field conditions. Interestingly, we saw few differences in species richness or microbiome community structure in mosquitoes caught in either trap. Co-occurrence and co-exclusion analysis identified 116 pairwise interactions substantially increasing the list of bacterial interactions observed in mosquitoes. Networks generated from the microbiome of *Ae. aegypti* often included highly interconnected hub bacteria. There were several instances where co-occurring bacteria co-excluded a third taxa, suggesting the existence of tripartite relationships. Several associations were observed in multiple species or in field and laboratory-reared mosquitoes indicating these associations are robust and not influenced by environmental or host factors. To demonstrate that microbial interactions can influence colonization of the host, we administered symbionts to *Ae. aegypti* larvae that either possessed or lacked their resident microbiota. We found that the presence of resident microbiota can inhibit colonization of particular bacterial taxa. Our results highlight that microbial interactions in mosquitoes are complex and influence microbiome composition.

## Introduction

The microbiome of mosquitoes can be highly variable, both within and between species, and is often dominated by relatively few genera (Boissière et al., 2012; Buck et al., 2016; Muturi et al., 2017; Osei-Poku et al., 2012; Wang et al., 2011). Understanding the factors that influence this variation is important as microbes drastically alter host biology. For mosquitoes, bacteria can affect a diverse number of traits including immunity, reproduction, survival, and vector competence (Hegde et al., 2015; Jupatanakul et al., 2014). These phenotypes have ramifications for the vectorial capacity of pathogens, and as such, microbial-based vector control strategies are under investigation to reduce the burden of arthropod-borne diseases (Bourtzis et al., 2014; Dennison et al., 2014; Saldaña et al., 2017). While our understanding of the contributing factors that affect the composition and abundance of the microbiome is expanding, there are still many unanswered questions regarding microbiome assembly and maintenance within mosquito hosts.

Exposure to environmental microbes is undoubtedly a major influence on the mosquito microbiome. These effects are particularly pronounced at the aquatic stage as larvae and pupae are immersed in water and can acquire bacteria by filter feeding. Indeed, several studies have shown the larval stages possess a similar microbiome as their larval water environment (Duguma et al., 2013; Gimonneau et al., 2014; Vázquez-martínez et al., 2009), and exposure to bacteria at these immature stages has implications for adult traits (Dickson et al., 2017). Furthermore, the larval habitat can influence the composition of the adult microbiome. Bacteria can be transstadially transmitted to the adult (Chen et al., 2015; Coon et al., 2014; Gonçalves et al., 2014; Jadin et al., 1966), and newly emerged adults are known to imbibe their larval water, which likely seeds the gut with microbiota (Lindh et al., 2008).

Host and bacterial genetics also contribute to microbiome composition and microbial abundance. Mosquitoes can maintain microbiome homeostasis by a variety of different mechanisms. Host pathways and processes known to influence microbiota in mosquitoes include immunity, amino acid metabolism, reactive oxygen species and calcium transport (Kumar et al., 2010; Pang et al., 2016; Short et al., 2017; Stathopoulos et al., 2014; Xiao et al., 2017; Zhao et al., 2017). Additionally, serial passaging of gut symbionts in mosquitoes selected for isolates that persist in the gut for longer periods of time (Dennison et al., 2016; Riehle et al., 2007), indicating that bacterial genetics is also important in shaping the microbiome.

Adult mosquito feeding behavior also has important implications for microbiome community structure. It is likely that bacteria can be acquired from the nectar of plants (Gusmão et al., 2007), and taking a blood meal alters the microbiome considerably. At 24 hours post-blood meal, the bacterial load in the gut drastically increases while species diversity decreases (Kumar et al., 2010; Oliveira et al., 2011; Terenius et al., 2012; Wang et al., 2011). Culture based assays show that bacterial loads revert to pre-blood fed levels 2-3 days after the blood meal (Demaio et al., 1996; Oliveira et al., 2011; Pumpuni et al., 1996), although other studies have seen high bacterial loads persist for some time and species richness not reverting to the original composition seen prior to the blood meal (Gusmão et al., 2010; Wang et al., 2011). Most of these studies either used laboratory-reared mosquitoes to examine culturable bacterial load, or relocated field mosquitoes to the laboratory for experimentation, and as such, there are few studies examining the effect of blood feeding on the microbiome community structure in field populations.

Recently, it has become evident that a further force affects microbiome composition in mosquitoes – interactions between the microbes themselves. These interactions were first highlighted with the discovery that *Wolbachia* and *Asaia* are antagonistic to one another, thereby affecting the vertical transmission of *Wolbachia* in *Anopheles* mosquitoes (Hughes et al., 2014; Rossi et al., 2015). Further comparisons exploiting 16S rRNA amplicon high throughput sequencing have identified interactions between *Wolbachia* and other microbes in both *Drosophila* and mosquitoes (Audsley et al., 2017b; Novakova et al., 2017; Simhadri et al., 2017; Ye et al., 2017; Zink et al., 2015). In addition to the specific interactions between *Wolbachia* and other bacterial taxa, pairwise negative and positive microbial interactions within bacteria or fungi, as well as cross-kingdom interactions (bacterial-fungal) have been reported in the La Crosse virus vectors, *Aedes triseriatus,* and *Ae. japonicus* (Muturi et al., 2016a). Taken together, these studies suggest that microbial interactions are important in dictating the composition and abundance of host-associated microbiota, yet it is unclear how ubiquitous and complex these interactions are within mosquitoes. 16S rRNA amplicon sequencing datasets have been analyzed to create microbial co-occurrence networks for several species and environments (Barberán et al., 2012; Chaffron et al., 2010; Faust and Raes, 2012; Faust et al., 2012; Goodrich et al., 2014), but these networks are lacking for mosquitoes and insects in general. These methods use presence/absence metrics, relative abundance, or both, to examine pairwise interactions to develop a network. Usually, interacting pairs of taxa are designated as having co-occurring or co-exclusionary relationships. Each method used for the identification of co-occurrence/co-exclusion networks has strengths and weaknesses in identifying particular patterns. CoNet (Faust and Raes, 2016), uses an ensemble approach that combines results from a collection of algorithms (Bray and Curtis, 1957; Cover and Thomas, 2012; Kullback and Leibler, 1951; Pearson, 1895; Sedgwick, 2014) using presence/absence and relative abundance data to identify statistically significant interactions. Interaction networks provide another methodology to examine the community structure of the microbiome of mosquitoes. Comparing microbiome networks generated from mosquitoes exposed to different conditions may provide insights into factors influencing microbiome structure in mosquitoes and identify pairwise interactions not affected by environmental conditions.

To expand our understanding of the forces that shape the bacterial microbiome of mosquitoes, we examined the microbial composition and community structure from three major mosquito arboviral vectors, *Ae. aegypti, Ae. albopictus,* and *Cx. quinquefasciatus,* collected from the field or reared under uniform insectary conditions. For the field collections, we utilized two trapping methods that primarily attract mosquitoes in different physiological states: host-or oviposition-seeking (Dennett et al., 2007; Figuerola et al., 2012; Maciel-de-Freitas et al., 2006; Reiter et al., 1986). Our sampling regime allowed us to examine how factors such as host species, environment, and physiological state in the field influenced the composition of the mosquito microbiome. We also compared the microbiome of mosquitoes that were naturally infected (Ae. *albopictus* and *Cx. quinquefasciatus)* and uninfected (Ae. *aegypti)* with *Wolbachia.* Furthermore, we developed microbial interaction networks to explore the complexity and nature of microbial interactions in mosquitoes. To demonstrate that microbial interactions influence host colonization, we infected *Ae. aegypti* larvae either possessing or lacking their native microbiota with a range of bacterial symbionts. Our results highlight the complexities of microbial networks in field collected *Ae. aegypti* mosquitoes and indicate the native microbiome induces colonization resistance to certain gut microbes.

## Material and Methods

### Mosquito collections and DNA extractions

Female *Ae. albopictus* and *Cx. quinquefasciatus* were collected from an abandoned tire repository in south-eastern Harris County, Houston, Texas, USA, while female *Ae. aegypti* were collected from a separate site in Houston (Figure S1). Further details describing the tire repository location were previously reported (Dennett et al., 2004). All mosquitoes were trapped over a 24 hour period with either the Biogents Sentinel (BG) or Harris County gravid (G) traps, which selectively collect host-seeking or gravid female mosquitoes, respectively (Dennett et al., 2007; Figuerola et al., 2012; Maciel-de-Freitas et al., 2006). Mosquito species were identified using morphological characteristics, surface sterilized (5 min in 70% ethanol followed by 3 washes in 1X PBS each for 5 min), and stored in ethanol at −20°C while awaiting DNA extraction. 5-7 day old adult sugar fed laboratory-colonized mosquitoes (*Ae. aegypti;* Galveston strain, *Ae. albopictus;* Galveston strain, and *Cx. quinquefasciatus;* Houston strain) were reared under conventional conditions and then processed in the same manner as field samples. All laboratory-reared mosquitoes were reared in the insectary at the University of Texas Medical Branch.

### High throughput sequencing and bioinformatics analysis

High-throughput sequencing of the bacterial 16S ribosomal RNA gene was performed using gDNA isolated from each sample. Sequencing libraries for each isolate were generated using universal 16S rRNA V3-V4 region primers (Klindworth et al., 2012) in accordance with Illumina 16S rRNA metagenomic sequencing library protocols. The samples were barcoded for multiplexing using Nextera XT Index Kit v2. Sequencing was performed on an Illumina MiSeq instrument using a MiSeq Reagent Kit v2 (500-cycles). The NCBI Bioproject accession number for the raw sequencing data reported here is PRJNA422599.

To identify the presence of known bacteria, sequences were analyzed using the CLC Genomics Workbench 8.0.1 Microbial Genomics Module (http://www.clcbio.com). Reads containing nucleotides below the quality threshold of 0.05 (using the modified Richard Mott algorithm) and those with two or more unknown nucleotides or sequencing adapters were trimmed out. All reads were trimmed to 264 bases for subsequent operational taxonomic unit (OTU) classification. Reference based OTU picking was performed using the SILVA SSU v119 97% database (Quast et al., 2013). Sequences present in more than one copy but not clustered to the database were placed into de novo OTUs (97% similarity) and aligned against the reference database with 80% similarity threshold to assign the “closest” taxonomical name where possible. Chimeras were removed from the dataset if the absolute crossover cost was 3 using a k-mer size of 6. Alpha diversity was measured using Shannon entropy (OTU level), rarefaction sampling without replacement, and with 100,000 replicates at each point. Beta diversity was calculated using the Bray-Curtis diversity measure (OTU level). PERmutational Multivariate ANalysis Of VAriance (PERMANOVA) analysis was used to measure effect size and significance on beta diversity for grouping variables (Anderson, 2014). The significance is obtained by a permutation test. For each assessment, a permutation of 99,999 was chosen. Differentially abundant bacteria (genus level, >0.1%) were identified using analysis of composition of microbiomes (ANCOM) (Mandal et al., 2015) with a significance level of P > 0.05, while values quantifying fold change were obtained using the log2 fold change formula (Quackenbush, 2002).

### Detection of complex interaction patterns

OTUs with read counts below 0.1% of total number of reads in all samples were excluded from analysis. The remaining OTUs were combined based on lowest common taxonomy assignments down to genus level, and relative abundance tables were generated by normalizing read counts against total number of reads in the original data. The resulting number of unique entries identified in samples was 33. Interactions (such as co-occurrence and co-exclusion) among these were identified using CoNet app (Faust and Raes, 2016) in Cytoscape (Shannon et al., 2003) using the following ensemble of methods: Pearson correlation (Pearson, 1895), Spearman correlation (Sedgwick, 2014), mutual information (Cover and Thomas, 2012), Bray-Curtis dissimilarity (Bray and Curtis, 1957) and Kullback-Leibler divergence (Kullback and Leibler, 1951). Statistical significance of each pair was tested using the row-shuffle randomization option and interactions that scored at the top and bottom 1% of 100 bootstraps were reported. Resulting statistically significant interactions were categorized by the software into three groups: co-presence, co-exclusion and unknown. Unknown interactions represent statistically significant patterns that cannot be clearly categorized as co-exclusion or co-occurrence. Since we could not ascribe an interaction pattern, the unknown interactions were excluded from the network. Resulting interaction networks were visualized using Cytoscape software (Shannon et al., 2003).

### Estimation of microbial load and screening for *Wolbachia* by PCR

Total bacterial load within each mosquito species or group was assessed by qPCR using gDNA as a template. qPCR was conducted using universal bacterial primers (Kumar et al., 2010) that amplified the bacterial 16S rRNA gene or primers that specifically amplified *Wolbachia* (Rao et al., 2006; Walker et al., 2009). Relative abundance was calculated by comparing the load of all bacteria or *Wolbachia* to a single copy mosquito gene (Calvitti et al., 2015; Isoe et al., 2011; Xia and Zwiebel, 2006). PCRs amplifying the 16S rRNA and *wsp* genes of *Wolbachia* were used to screen for *Wolbachia* in field caught *Ae. aegypti* (Baldo et al., 2006; O’Neill et al., 1992; Werren and Windsor, 2000; Zhou et al., 1998), and nematode specific primers were used to screen for nematode infections (Casiraghi et al., 2004).

### Re-infection of bacteria into mosquito larvae

*Aedes aegypti* gnotobiotic larvae were generated as previously described (Coon et al., 2014). To synchronize hatching, sterile eggs were transferred to a conical flask and placed under a vacuum for 45 min. To verify sterility, larval water was plated on non-selective LB agar plates and reared under sterile conditions. L1 larvae grown without bacteria have slow growth rates and do not reach pupation (Coon et al., 2014). Forty-five L1 larvae were transferred to a T75 tissue culture flask and inoculated with 1×10^7^ CFU/ml of transgenic symbionts possessing the plasmid the pRAM18dRGA-mCherry that was derived from pRAM18dRGA (Burkhardt et al., 2011). Bacterial cultures were quantified with a spectrophotometer (DeNovix DS-11, DeNovix) and validated by plating to determine CFU. For conventional rearing of mosquitoes, eggs (non-sterilized) were vacuum hatched and grown under non-aseptic conditions in a T75 tissue culture flask supplemented with transgenic symbionts at the concentration of 1×10^7^ CFU/ml. To feed mosquitoes, ground fish food pellets were sterilized by autoclaving, and mixed with sterile water. The equivalent of 6 mg of fish food was fed to both gnotobiotic and conventionally reared mosquitoes every second day. To quantify the bacterial load of symbionts, surface sterilized L4 larvae were homogenized and plated on a selective media (50 μg/ml Kanamycin) on which only transgenic symbionts grew (pRAM18dRGA-mCherry induces resistance to Kanamycin). After incubation at 30°C or 37°C (depending on symbiont) for 2-3 days, colonies (expressing mCherry fluorescent protein) were counted. All colonies observed on the kanamycin plate were confirmed to have mCherry fluorescence.

## Results

### Microbiome Diversity

We sequenced amplicons of the V3/V4 region of the 16S rRNA gene from whole individual adult female mosquitoes either collected from the field or reared in the laboratory. In total, we sequenced 130 adult mosquitoes obtaining 10,668,291 reads (sample size per group and species is reported in Table S1). After quality filtering, 7,051,256 reads were assigned to OTUs at 97% identity threshold and on average, there were 54,240 reads per mosquito sample. Rarefaction curve analysis indicated that our sequencing depth was sufficient to observe all OTUs in mosquito samples (Figure S2). We identified a total of 4,419 bacterial OTUs in the three mosquito species, but only 58 were present at an infection frequency of over 0.1% within the dataset (Table S2). When abundant microbes were classified at higher taxonomic levels, our analysis found 22 families, with Enterobacteriaceae being the most common when disregarding *Wolbachia.* Bacteria found in mosquitoes were classified into five phyla with bacteria in the phylum Proteobacteria most prevalent in the microbiome, which is consistent with previous studies (Audsley et al., 2017a; David et al., 2016; Muturi et al., 2016b; Osei-Poku et al., 2012), although other reports indicated Bacteroidetes or Acinetobacter phyla can be a major component of the microbiome (Coon et al., 2014; 2016b; Minard et al., 2014; 2015).

When examining the sequencing data at the genus level, the microbiomes of *Cx. quinquefasciatus* and *Ae. albopictus* were dominated by the endosymbiont *Wolbachia* with 87 and 81 % of total reads, respectively (Figure S3, Table S2). While *Wolbachia* accounted for many of the reads, rarefaction analysis indicated our sampling depth was sufficient to identify rare OTUs. Other highly abundant genera in field-collected mosquitoes included *Halomonas, Shewanella,* and *Asaia* in *Cx. quinquefasciatus, Halomonas, Pseudomonas,* and *Zymobacter* in *Ae. albopictus,* and *Pseudomonas, Zymobacter, Tatumella,* and *Enterobacter* in *Ae. aegypti* (Figure S3, Table S2). Similar to a recent finding (Coon et al., 2016b), we found a small number of *Wolbachia* reads in a few *Ae. aegypti* individuals collected from the field in G traps. It is thought that *Ae. aegypti* are naturally uninfected by *Wolbachia* (Iturbe-Ormaetxe et al., 2011), although Coon *et al.* (2016) suggested that some populations may be infected. However, samples that contained *Wolbachia* reads from our sequencing data could not be independently validated by PCR using several *Wolbachia* genes *(wsp* and MLST genes) commonly used to screen for the bacterium, nor were they PCR positive for filarial nematode infection (that carry *Wolbachia)* when amplifying with primers that detect nematode DNA.

Given the above finding and since microbiome sequencing can be susceptible to contamination (Pollock et al., 2018; Tourlousse et al., 2017), we examined our data for other possible contamination signatures. While we could not find conclusive evidence of contamination, in this process we observed a possible batch effect for a specific bacterium in field-collected *Ae. aegypti.* These samples were extracted in two batches (see Table S1) and the field-collected samples from the latter extraction were found to have higher loads of *Burkholderia* compared to those extracted in the first batch. However, all laboratory-reared mosquitoes extracted during this second extraction did not contain high levels of *Burkholderia,* indicating the prevalence of this microbe in the field-collected samples was not likely due to laboratory contamination. Despite this occurrence, the dataset is suitable for constructing networks as this analysis examines pairwise interactions, and as such, other interacting pairs will not be influenced by *Burkholderia.* Future studies should consider the use of reagent-only controls and spike controls to help determine if cross-contamination of samples occurs in the sequencing process (Pollock et al., 2018; Tourlousse et al., 2017), although it is questionable if these controls would have been of assistance in this case given that laboratory-reared mosquitoes did not have elevated *Burkholderia* reads.

The Shannon diversity index was used to estimate the species richness in mosquitoes (Shannon, 2001). There were significant differences in diversity between the microbial communities of *Aedes* mosquitoes compared to *Cx. quinquefasciatus* when compared at the OTU level (Figure 1; Kruskal-Wallis P<0.0001). When comparing groups within *Ae. aegypti,* field collected mosquitoes from either the BG or G trap had a significantly greater Shannon diversity index compared to the laboratory mosquitoes. No differences were seen between groups in *Ae. albopictus,* while *Cx. quinquefasciatus* caught in the G trap had a significantly lower Shannon diversity index compared to laboratory-reared mosquitoes. Across all species, we found no significant differences in species richness between mosquitoes caught in the BG and G traps.

**Figure. 1.**
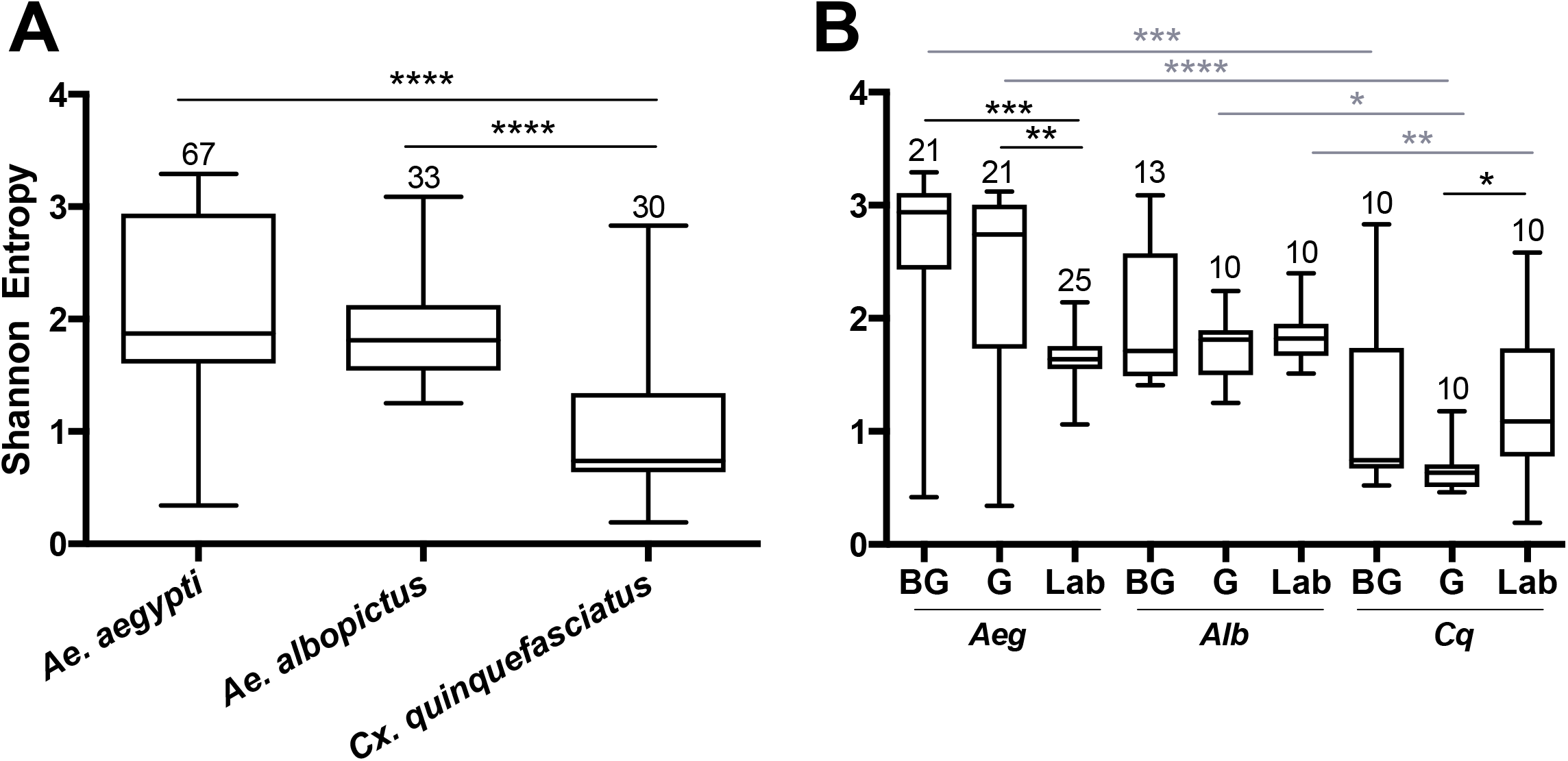
Shannon diversity indices at the OTU level for all mosquito species (A) or for each group within a species (B). A Kruskal-Wallis test with a Dunn’s multiple comparison test was used to determine significance (* P< 0.05, ** P<0.01, *** P<0.001, **** p<0.0001) within a species (black) or group (grey). Abbreviations: *Aeg* – *Ae. aegypti, Alb – Ae. albopictus, Cq – Cx. quinquefasciatus.* Bars on the box plots show maximum to minimum range. Sample size for species and group is indicated by the numbers above the box-plots and in Table S1.

Since both *Ae. albopictus* and *Cx. quinquefasciatus* were heavily infected with *Wolbachia,* we examined alpha diversity (OTU level) in these mosquitoes when this endosymbiont was computationally excluded from the dataset (Figure S4). In all cases, we observed an increase in Shannon diversity when *Wolbachia* was excluded. This was significant when analyzed by species (Ae. *albopictus* P < 0.0001; *Cx. quinquefasciatus* P < 0.0001), and for all groups with the exception of *Ae. albopictus* caught in the G traps (Ae. *albopictus:* BG P < 0.005, Lab P < 0.02; *Cx. quinquefasciatus:* BG P < 0.04, G P < 0.001, Laboratory P < 0.01). No significant differences were seen when comparing the BG to G groups after removal of *Wolbachia (Ae. albopictus:* P = 0.23; *Cx. quinquefasciatus:* P = 0.57).

### Factors that influence microbiome community structure

We examined how the environmental, physiological state in the field, and host species affected the bacterial community structure using beta diversity analysis by comparing the microbiomes of the three mosquito species or groups. Within each group, distinct microbiome clustering patterns were observed between the three mosquito species and all pairwise comparisons were significantly different (Figure 2; PERMANOVA with Bray-Curtis distance comparison P<0.05). In general, the microbiome of *Ae. aegypti* was more divergent compared to the microbiomes of *Cx. quinquefasciatus* and *Ae. albopictus* regardless of origin (Figure 2A). When *Wolbachia* was computationally excluded from the analysis (Figure 2A), the microbiomes of *Cx. quinquefasciatus* and *Ae. albopictus* became more divergent in the laboratory, but were not significantly different when considering mosquitoes caught in the BG trap (P=0.24). Interestingly, there was tight clustering of samples from *Cx. quinquefasciatus* and *Ae. albopictus* caught in the G trap (Figures 2A), although these two groups were significantly different (P=0.00001). When *Wolbachia* was removed, this clustering became more divergent, yet still was significantly different (P=0.0052) (Figure 2A).

**Figure. 2.**
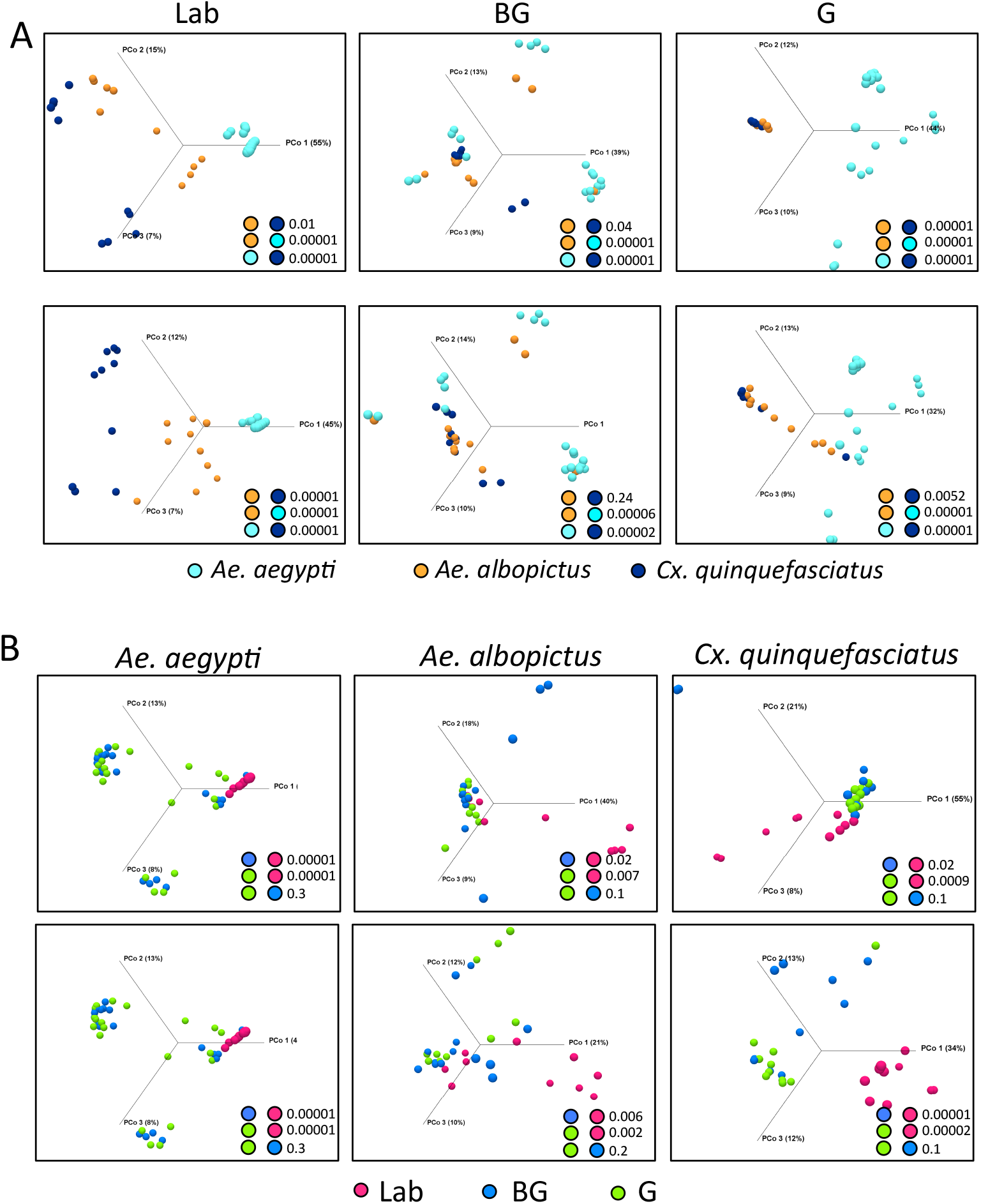
Principal Coordinates Analysis (OTU level) using Bray-Curtis dissimilarity, comparing identified OTUs within a group (A) or species (B). PERMANOVA significance values for pairwise comparison are reported in the lower right corner for each analysis. For A and B, upper plots include *Wolbachia* while *Wolbachia* has been computationally excluded in the lower plots.

To determine how the environment and physiological state influences microbial composition we compared the microbiome of laboratory-reared and field-caught (BG or G) individuals within each mosquito species. For all species, laboratory-reared mosquitoes had a significantly distinct microbiota compared to their field counterparts (Figure 2B). This was most pronounced in *Ae. aegypti* but less distinct for *Ae. albopictus* and *Cx. quinquefasciatus.* No significant differences were observed in the microbiome community structure of mosquitoes caught in the BG or G traps for any of the three mosquito species. No differences were observed when these microbiomes were analyzed with *Wolbachia* computationally excluded (Figure 2B).

### Common and differentially abundant bacterial between and within mosquito species

We examined our data for bacterial genera that were unique to or shared between mosquito species. The majority of bacteria were common between species with the notable exception that *Ae. aegypti* caught in the G trap possessed 12 genera not present in the other two mosquito species (Figure 3A, Table S3). Similarly, when comparing within a species between groups, most bacteria were common to all groups (Laboratory, BG, G). To detect bacteria that may contribute to the observed differences in microbiome community structure of a particular mosquito species, we completed pairwise comparisons to identify bacteria that were differentially abundant (Figure 3B). These comparisons were completed within each group (Laboratory, BG or G). The largest differences in the microbiome were seen when comparing *Ae. aegypti* to the other two species, which agreed with our beta diversity analysis findings. Several bacterial taxa were differentially abundant between species regardless of group (Laboratory, BG and G traps), suggesting environmental factors are not greatly influential on these specific host-microbe associations. For example, *Aeromonas, Serratia, Shewanella* and *Wolbachia* were less abundant in *Ae. aegpyti* compared to both *Ae. albopictus* and *Cx. quinquefasciatus* regardless of environmental conditions. When comparing differential abundance within a group, we were able to find infection gradients of specific microbes across species. The two examples of these infection clines were for *Serratia* and *Aeromonas* in laboratory-reared mosquitoes. These bacteria heavily infected *Cx. quinquefasciatus,* had moderate infection densities in *Ae. albopictus* and poorly infected or were absent from *Ae. aegypti,* despite the fact these three mosquito species were reared under common environmental laboratory conditions (Figure 3B and Figure S5).

**Figure. 3.**
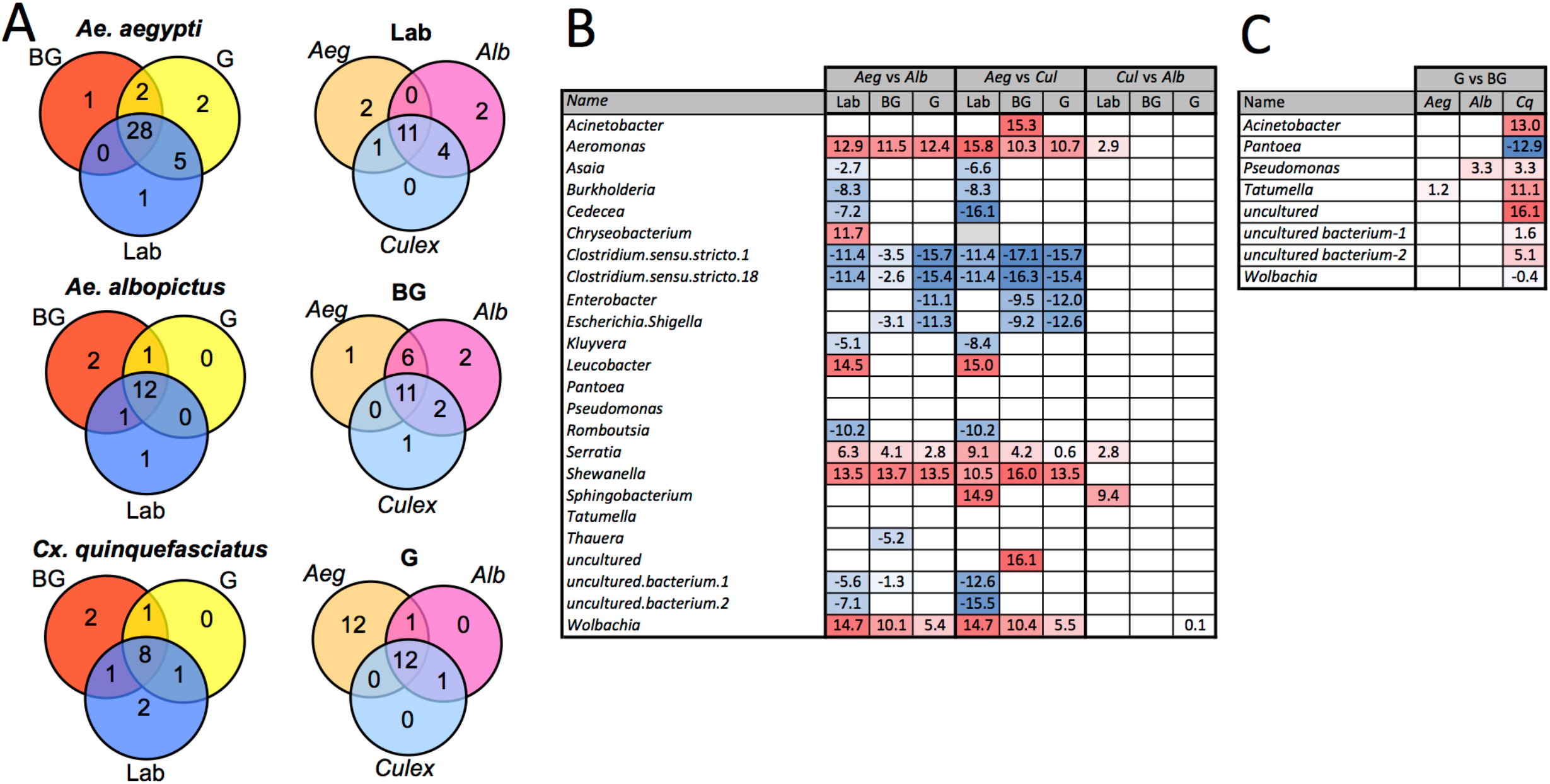
Common and differentially abundant bacteria (genus level) within mosquitoes. Venn diagram showing number of common bacterial genera between mosquito groups and species (A). Pairwise comparison of bacterial density between each mosquito species within each group (B). Pairwise comparison of bacterial density for mosquito caught in the G trap compared the BG trap (Ae. *aegypti* – *Aeg, Ae. albopictus* – *Alb, Cx. quinquefasciatus* – Cq.) (C). Log_2_ values indicated fold change in bacterial density. List of common taxa for each species and group are presented in Table S3.

Several studies have reported that blood feeding alters the species richness in mosquito guts (Kumar et al., 2010; Oliveira et al., 2011; Terenius et al., 2012; Wang et al., 2011). While at the community level, the microbiomes of G and BG were not significantly different (Figure 2), we did find specific bacteria that were differentially abundant between these groups (Figure 3C). These changes were mainly seen in *Cx. quinquefasciatus.* Of the known bacteria, the greatest changes were seen in *Acinetobacter, Tatumella,* and *Pantoea,* with the former two being more abundant in the mosquitoes caught in BG traps, while the latter was more abundant in mosquitoes caught in the G trap.

### Total bacterial load in mosquitoes

While high throughput sequencing allows characterization of the composition of the microbiota, it only provides a relative measure of bacterial density (Gloor et al., 2017). Therefore, to obtain an estimate of the total bacterial load in each vector species, we completed qPCR on mosquitoes with universal eubacterial primers that broadly amplify bacterial species (Kumar et al., 2010). *Culex* mosquitoes were seen to have a higher total bacterial load when compared to either of the two *Aedes* species (Figure 4; Kruskal Wallis P<0.0001). When comparing within a species between groups, we found that laboratory-reared *Ae. aegypti* had significantly greater load than those caught in the field (Kruskal Wallis P<0.0001). This was also the case for *Cx. quinquefasciatus* (Kruskal Wallis P<0.0001) while no significant differences were seen between *Ae. albopictus* groups. As both *Ae. albopictus* and *Cx. quinquefasciatus* are infected by *Wolbachia,* we also quantified *Wolbachia* by qPCR to determine its relative density in proportion to the total bacterial load of mosquitoes. While it is inappropriate to statistically compare amplicons that have different amplification efficiencies, it is evident that *Wolbachia* comprises a major component of the microbiome in *Ae. albopictus* and *Cx. quinquefasciatus,* which corroborates the high throughput sequencing data.

**Figure. 4.**
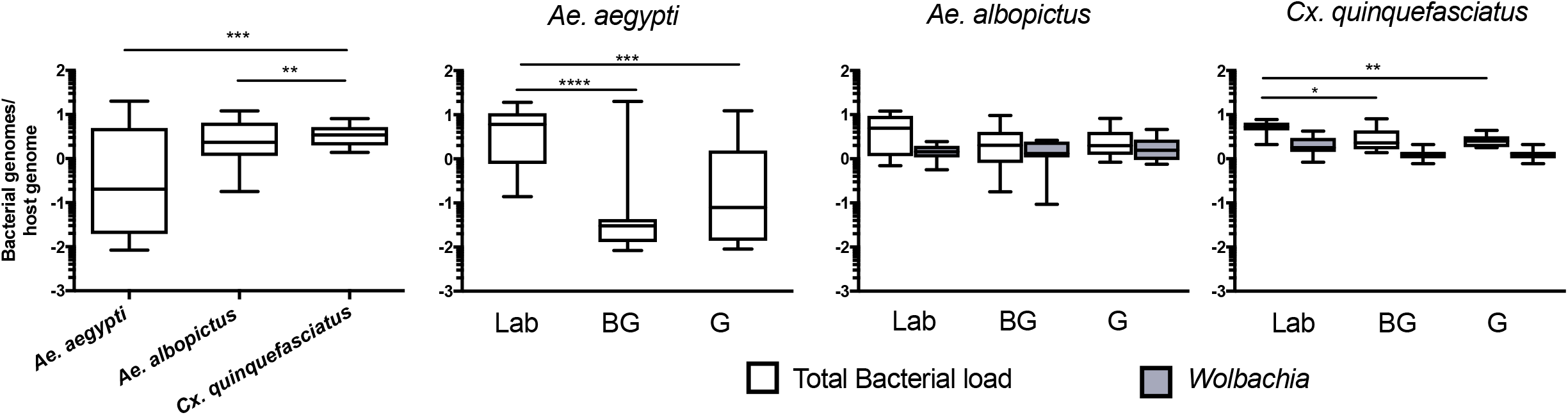
Total bacterial load of mosquitoes. Comparison of bacterial load for each species (left) or within each group for each mosquito species (right). Bacterial load is represented as a ratio between 16S rRNA gene copies to S7 copies (Ae. *aegypti* and *Cx. quinquefasciatus)* or actin (Ae. *albopictus)* genes. The density of *Wolbachia* was estimated for *Ae. albopictus (Wolbachia 16s:Actin)* and *Cx. quinquefasciatus (ftsZ:S7).* A Kruskal-Wallis test with a Dunn’s multiple comparison test was used to determine significance (* P < 0.05, ** P<0.01, *** P<0.001, **** p<0.0001) of total bacterial loads within a species. Bars on the box plots show maximum to minimum range. Sample size for all species and groups is described in Table S1.

### Microbial interaction networks within mosquitoes

The 16S rRNA sequencing data can be analyzed to create microbial interaction networks, providing information on potential interaction patterns of microbes such as co-occurrence and co-exclusion. We created network maps of bacterial interactions using a variety of models that use presence/absence and relative abundance data to identify pairwise relationships (Faust and Raes, 2016). In general, we saw that interaction networks from *Ae. aegypti* were complex, in that they had more nodes and connections compared to networks from *Ae. albopictus and Cx. quinquefasciatus* (Figure 5, Table S4). For all mosquito species, both co-occurrence and co-exclusion interaction patterns were observed in all networks (examples of these patterns are shown in Figure S6). We were able to identify taxa, or groups of bacteria, within these interactions that appear to be important to the overall structure of the network. For example, in the *Ae. aegypti* networks from field collected mosquitoes, *Enterobacter* and *Pseudomonas* are highly interconnected species having between 6 and 15 interactions in these networks. In the laboratory-reared *Ae. aegypti,* an *Enterobacteriaceae* had several interactions with other bacteria. Three-way interactions were also seen in many of the networks, and often these involved *Pseudomonas and Enterobacter.* Examples of tripartite co-occurrences interactions in *Ae. aegypti* networks include *Pseudomonas, Asaia* and a *Clostridium* isolate in laboratory-reared mosquitoes, *Pseudomonas, Serratia,* and *Enterobacter* and *Pseudomonas, Acidovorax,* and *Enterobacter* in the G and BG groups, respectively. Common co-exclusionary interactions were found in the networks generated from field-collected mosquitoes such as *Pseudomonas* co-excluding *Pantoea* and *Tatumella. Burkholderia* co-excluded *Enterobacter, Acidovorax* and *Escherichia-Shigella,* however, these *Burkholderia* interactions could possibly be an artifact due to extraction batch variation.

**Figure. 5.**
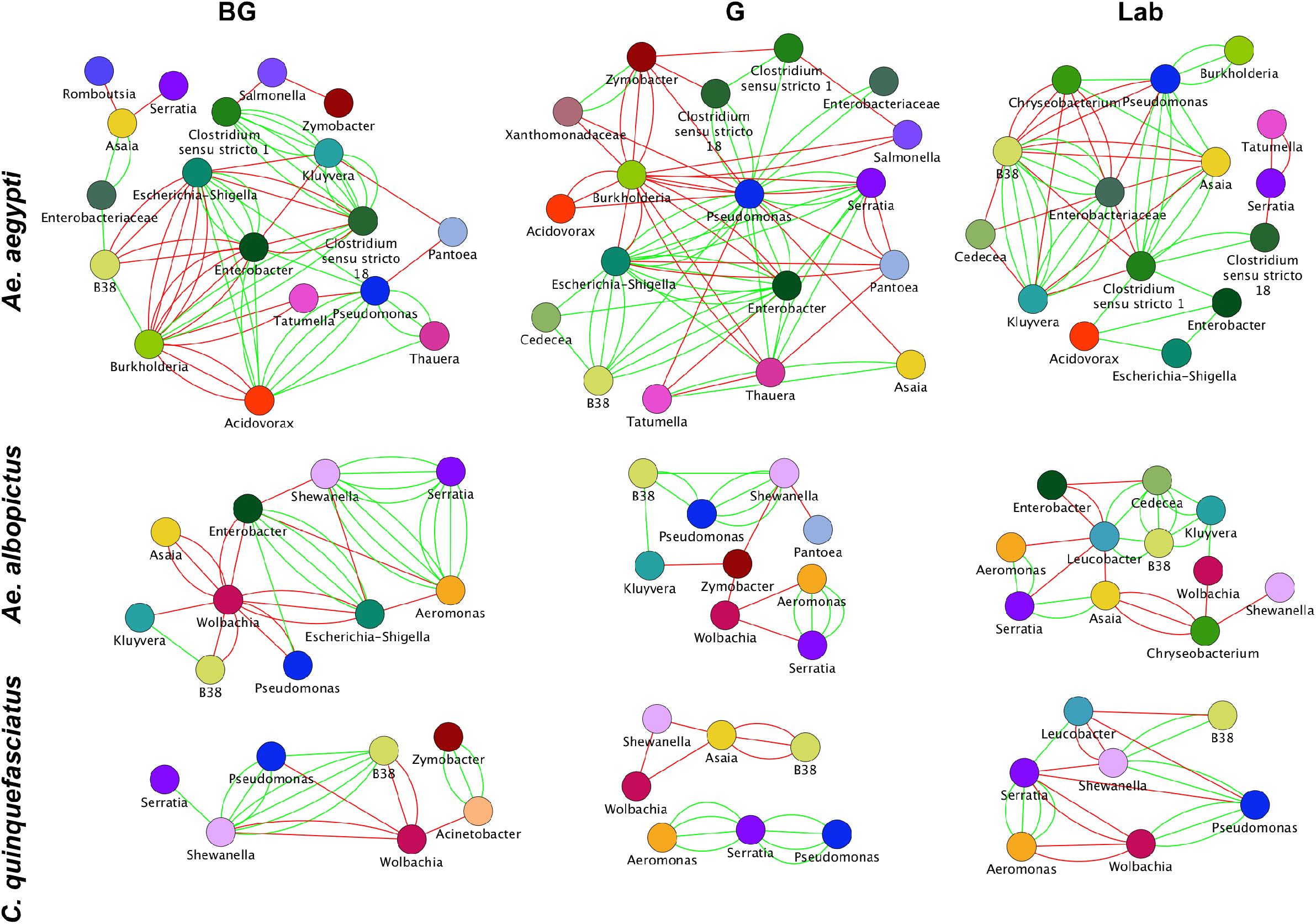
Microbial interaction networks for mosquitoes. Interaction networks were built using CoNet. Node colors represent unique taxonomy identifiers. Red edges represent co-exclusion/negative correlation, green edges represent co-occurrence/positive correlation interactions between relative abundance profiles. Multiple edges connecting the same nodes indicate significance from more than one metric (Bray-Curtis dissimilarity, Kullback-Leibler divergence, mutual information, Spearman correlation, and Pearson correlation). Undetermined interactions have been removed from the network.

*Wolbachia* was a highly interconnected taxon in the interaction networks generated from *Ae. albopictus* and *Cx. quinquefasciatus* mosquitoes, and often had co-exclusionary relationship with other bacteria. In BG-collected *Ae. albopictus, Wolbachia* co-excluded six other bacteria including *Asaia* and *Pseudomonas.* In other groups, *Wolbachia* was seen to repeatedly exclude *Aeromonas, Serratia* and *Shewanella.* The three-way interaction of *Wolbachia* co-excluding the co-occurring *Aeromonas* and *Serratia* was observed in both *Ae. albopictus and Cx. quinquefasciatus.* The *Aeromonas* and *Serratia* co-occurrence pattern appears highly robust and independent of environmental factors as this interaction was observed in five of the six *Ae. albopictus* and *Cx. quinquefasciatus* groups.

### Artificial infection of symbionts in germ-free or septic mosquitoes

The microbial interaction networks highlight the multifarious interactions that occur within mosquito systems. We undertook preliminary validation experiments using *Ae. aegypti* larvae to further demonstrate that microbial interactions can influence microbiome composition and abundance. To this end, we exploited the recently developed gnotobiotic rearing approach where mosquito larvae can be infected with a single bacterial taxon (Coon et al., 2014). We compared the density and prevalence of artificially infected symbionts in gnotobiotic lines (infected with the single symbiont) compared to conventionally reared septic mosquitoes (that possessed their resident microbiota) (Figure 6). We completed this with six bacteria isolated from *Aedes* mosquitoes. Three of the bacteria were isolated from *Ae. aegypti (Pantoea, Cedecea [Cedecea-aeg], Asaia)* while another three were isolated from *Ae. albopictus (Serratia, Cedecea [Cedecea-alb],* and *Enterobacter).* Three bacteria *(Serratia;* Mann Whitney P<0.0001, *Cedecea-alb;* Mann Whitney P<0.0001, and *Cedecea-aeg;* Mann Whitney P<0.0001) were observed to significantly infect *Ae. aegypti* mosquitoes at higher densities when inoculated into axenic rather than non-axenic mosquitoes. The prevalence (number of individuals infected) of *Serratia* (Fisher’s exact test, P<0.0001) and *Cedecea-alb* (Fisher’s exact test, P<0.0002) was also significantly higher in gnotobiotic compared to conventionally reared mosquitoes. No change in either the prevalence or density of infection was seen for *Enterobacter, Pantoea* or *Asaia.* Taken together, these data indicate that microbial interactions influence colonization and infection dynamics of specific bacterial species within mosquitoes.

**Figure. 6.**
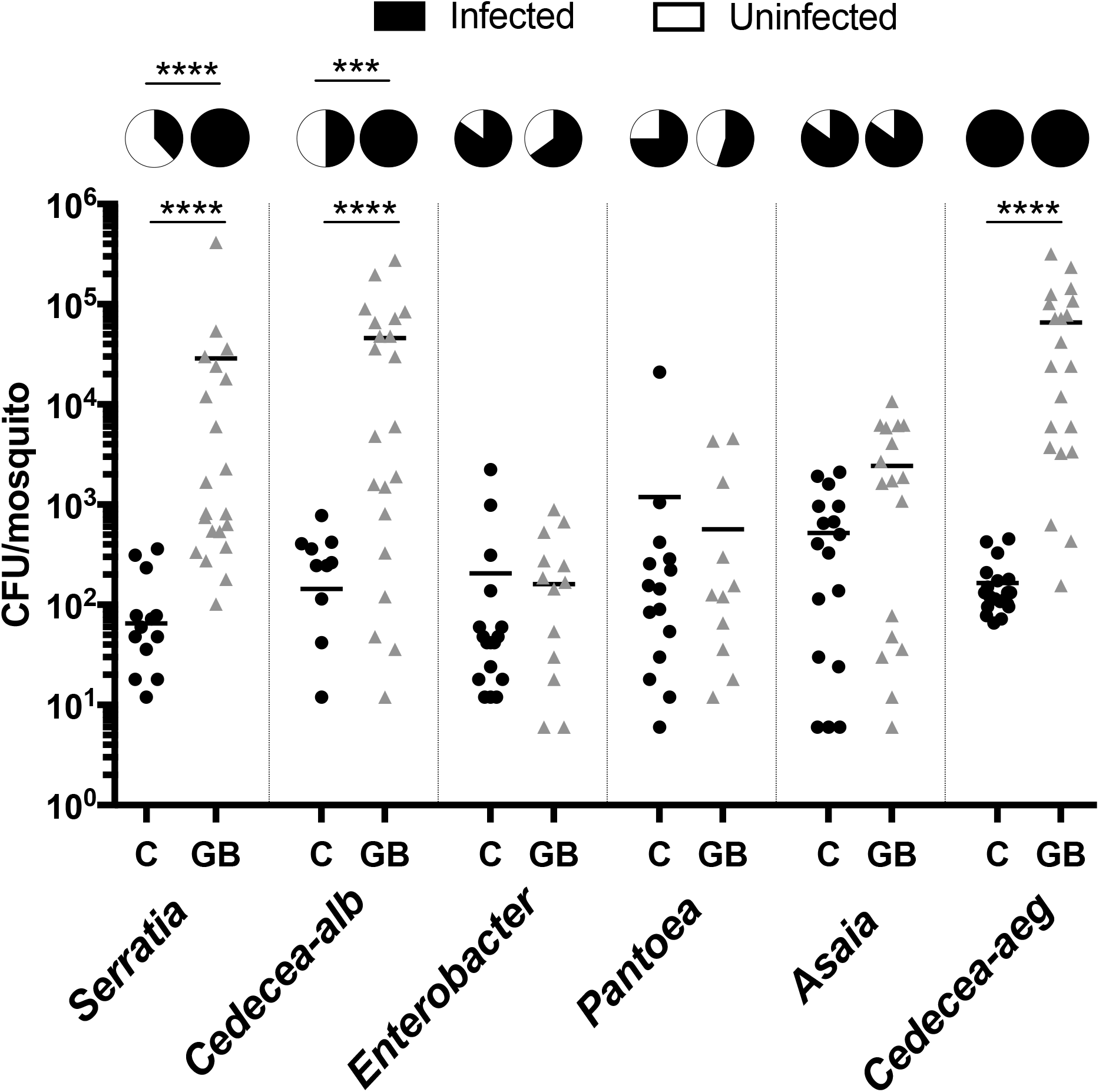
Infection density and prevalence of bacteria inoculated into conventionally reared or axenic mosquitoes. As such, gnotobiotic reared lines (GB) only possessed the inoculated bacteria while conventionally reared lines (C) possess their native microbiota in addition to the inoculated strain. Inoculated bacteria possessed a plasmid expressing antibiotic resistance and mCherry fluorescent protein. The bacterial load was quantified by counting mCherry colonies on selective plates. *Serratia, Cedecea-alb* and *Enterobacter* were isolated from *Ae. albopictus* (Galveston) while *Pantoae, Asaia* and *Cedecea-aeg* were isolated from *Ae. aegypti* (Galveston). A Mann-Whitney test with a Dunn’s multiple comparison test was used to determine significance (**** is P<0.0001). Pie charts indicate prevalence of infection (Fisher’s exact test, **** P<0.0001, *** P<0.0002).

## Discussion

It is clear that complex factors combine to shape the microbiome of an organism. To further increase our understanding of factors that affect the microbiome of mosquitoes, we sequenced the microbiome of laboratory-reared and field-caught adult mosquitoes exploiting traps that attract host- or oviposition-seeking individuals. Anautogenous mosquitoes require a blood meal to provide the necessary nutrition for egg development and this dramatic influx of blood has been shown to substantially alter the microbiome of laboratory-reared mosquitoes or those caught in the field and blood-fed in the laboratory (Kumar et al., 2010; Oliveira et al., 2011; Terenius et al., 2012; Wang et al., 2011). Intriguingly, we saw few differences in the microbiome between host- and oviposition-seeking mosquitoes. Across all species, there were no significant differences in alpha or beta diversity for mosquitoes caught in either trap. Similarly, the total bacterial load was similar between mosquitoes caught in the BG and G traps, and this was consistent across all three species. However, when comparing beta diversity between traps, we did see less variation in the groupings of individuals from both *Ae. albopictus* and *Cx. quinquefasciatus* in the G trap compared to the BG trap. Overall, our results are in contrast to studies using lab-reared mosquitoes or mosquitoes caught in the field and blood-fed in the laboratory that indicate increased bacterial load, but decreased diversity over a 24-48 hour window following a blood meal (Kumar et al., 2010; Oliveira et al., 2011; Terenius et al., 2012; Wang et al., 2011).

Several factors could explain the differences between our results examining field-collected samples and those of previous studies (Kumar et al., 2010; Oliveira et al., 2011; Terenius et al., 2012; Wang et al., 2011). First, mosquitoes often take multiple blood meals, particularly *Ae. aegypti* (Scott et al., 1993). In our collections, the post-blood feeding history of mosquitoes is unknown, and it is possible that mosquitoes caught in either trap may have had a blood meal (or several) prior to being caught. The body of work examining the influence of blood feeding on the microbiome has only examined the effect of a single blood meal, not several, and while it appears the microbiome reverts to a pre-blood fed microbiome several days post-blood meal, it is unknown how quickly this occurs in the field. In field collected *Anopheles* mosquitoes reared in the laboratory, reversion to a Bacteroidetes-dominated microbiome, which was the dominant phylum seen in sugar fed mosquitoes, was seen 4 days post-blood meal (Wang *et al.* 2011). In our samples collected from G traps, females may have gone several days without a blood meal before finding a suitable oviposition site (i.e., the trap), possibly even relying on nectar-based food sources for sustenance. It is also possible that mosquitoes in search of an oviposition site may have never taken a blood meal. While most mosquitoes usually require a blood meal to develop eggs, autogeny has been reported in these species (Ariani et al., 2015; Chambers and Klowden, 1994; Olejnícek and Gelbic, 2000). Autogeny rates, which are variable and depend on temperature and nutrition, have been reported to range from 3 – 34% for *Ae. aegypti,* around 5% for *Ae. albopictus,* and up to 87% in *Culex* mosquitoes (Ariani et al., 2015; Chambers and Klowden, 1994; Olejnícek and Gelbic, 2000; Trpis, 1977). Little is known regarding the influence of microbiota and autogeny, although in the autogenous mosquito *Ae. atropalpus,* specific gut taxa have been shown to influence egg production (Coon et al., 2016a). Finally, while reports indicate that BG and G traps preferential catch mosquitoes in different physiological states (Dennett et al., 2007; Figuerola et al., 2012; Maciel-de-Freitas et al., 2006; Reiter et al., 1986), we did not explicitly examine if females were gravid or not. While we contemplated dissecting mosquitoes to examine their parity, we chose not to, as it would increase the potential for contamination of the samples. Although in our work we did not see dramatic differences in the mosquitoes caught in BG or G traps, the earlier points highlight the challenges in undertaking studies on field-derived samples and could explain the disparity between our results and those from studies undertaken in laboratory settings.

Since both *Ae. albopictus* and *Cx. quinquefasciatus* are heavily infected with *Wolbachia,* we analyzed our data with and without this endosymbiont to garner a better understanding of the other bacterial microbiota in these mosquitoes. Our high throughput sequencing and qPCR results demonstrated that *Wolbachia* was the most abundant bacterium in *Ae. albopictus* and *Cx. quinquefasciatus.* In spite of this, we obtained sufficient sequencing depth to identify other bacterial taxa – a common challenge when characterizing Wolbachia-infected species using amplicon sequencing (Minard et al., 2014). Removing *Wolbachia* from our analysis increased the Shannon diversity index, indicating the remaining microbiota within these mosquitoes is relatively even. For beta diversity, we found that our results were mixed and dependent on group. When comparing within a group, removing *Wolbachia* made the microbiomes of laboratory-reared *Ae. albopictus* and *Cx. quinquefasciatus* more divergent while the microbiomes of field mosquitoes tended to be less different. These findings are consistent with a study by Novakova *et al* (2016) that found the removal of *Wolbachia* from their analysis led to less distinct differences for field-collected mosquitoes. When comparing groups within a species, exclusion of *Wolbachia* made the comparison of field and laboratory mosquitoes more distinct, likely due to differences in the gut-associated microbiota. Similar to a recently published study (Coon et al., 2016b), we found a small number of *Wolbachia* reads in *Ae. aegypti* mosquitoes collected in G traps. However, we could not confirm the presence of the bacteria with conventional PCR-based approaches, suggesting these results could be a sequencing or laboratory artifact. Should populations of *Ae. aegypti* be naturally infected with *Wolbachia,* this could have important ramifications for biological control strategies being implemented into the field (Bourtzis et al., 2014; Flores and O’Neill, 2018; Hughes and Rasgon, 2014), and as such, further research in this area is warranted.

Similar to findings in other mosquito species, many bacteria were shared between different mosquito species, and laboratory-reared mosquitoes were seen to have a divergent microbiome from their field counterparts (Boissière et al., 2012; Coon et al., 2016b). These common taxa, particularly those that infect at high abundance, could be candidate bacteria for consideration in novel pan-mosquito microbial control strategies as they would likely be compatible for all three vector species (Saldaña et al., 2017). When focusing specifically on bacterial titers, few bacteria were seen to be significantly different between mosquitoes caught in either traps. In *Culex* mosquitoes, *Pantoea* was more abundant in individuals caught in the G trap. Similar to this finding, *Pantoea* has been found to increase in abundance after a blood meal in *Anopheles* mosquitoes (Wang et al., 2012). When considering pairwise comparisons between species, there were more differentially abundant genera when comparing *Ae. aegypti* to the other two species, which explains the greater divergence of the *Ae. aegypti* microbiome to the microbiome of the other two species. The *Ae. aegypti* samples were collected from a different location than the other two species, thus, it is possible that environmental factors could explain these differences. However, we also saw common changes that were consistent regardless of groups (BG, G and Laboratory), such as *Aeromonas*, *Clostridium, Serratia, Shewanella* and *Wolbachia,* indicating these bacteria were not greatly influenced by the environment and that other factors affected their presence in the particular mosquito species. Notably, we found that in laboratory-reared mosquitoes, *Culex* harbored significantly higher titers of *Serratia* and *Aeromonas,* compared to *Ae. albopictus,* while these bacteria were at low abundance or absent from *Ae. aegypti.* These results suggest there are host and/or bacterial related factors that make this particular strain of *Ae. aegypti* (Galveston) inhospitable for *Serratia* and *Aeromonas* as all three mosquito species were subjected to similar uniform environmental conditions when reared in the insectary.

Current evidence of microbial interactions within mosquitoes is mainly limited to interactions between *Wolbachia* and other microbiota (Audsley et al., 2017a; Hughes et al., 2014; Rossi et al., 2015; Zink et al., 2015), or between vertically transmitted symbionts in other arthropod systems (Goto et al., 2006; Kondo et al., 2005; Macaluso et al., 2002; Rock et al., 2017). As such, our understanding is generally restricted to inherited symbionts and we have a poor understanding of the scale of interactions between microbes that are associated with insect guts. To address this, we created microbial interaction networks to identify pairwise co-occurrence and co-exclusion patterns. To avoid spurious interactions, which could be due to the presence or absence of a microbe in one environmental condition but not another, we limited our network analysis to within a group for each particular species. Our analysis identified 116 co-occurrence or co-exclusion interactions, substantially increasing the number of bacterial interactions observed in mosquitoes. Bacterial interaction networks generated from *Ae. aegypti* mosquitoes were more complex than *Ae. albopictus* or *Cx. quinquefasciatus* in that they had more nodes and connections. Species richness may explain the differences observed in network structure, as in general, the more complex networks had a greater number of OTUs. Other factors that may have influenced the identification of interacting bacteria are the presence of *Wolbachia* in *Ae. albopictus* and *Cx. quinquefasciatus,* as well as the differences in sample size between mosquito species. Further studies are warranted to determine why some networks are highly interconnected while others are not.

Interestingly, in the more complex networks, we saw evidence of hub microbial taxa that were highly interconnected. *Pseudomonas* and bacteria within the *Enterobacteriacaea* appear to be important hub taxa. Some of the interactions observed here have been previously reported in *Ae. triseriatus* and *Ae. japonicus* (Muturi et al., 2016a), including negative interactions between bacteria within the *Burkholderiaceae* and *Pseudomonas* and *Acinetobacter* as well as *Asaia-Enterobacter* and *Asaia-Ralstonia* interactions. The majority of interactions reported in *Ae. triseriatus* and *Ae. japonicus* were negative (Muturi et al., 2016a), whereas here we see a mix of both co-occurrence and co exclusion patterns. Specific hub microbes that are strongly interconnected have been found in plant microbiomes and these taxa have a profound effect on overall microbiome structure (Agler et al., 2016). In our data, we also saw three-way interactions. Often these were relationships were co-occurring, or were formed by two co-occurring bacteria both co-exclude another bacterium, suggesting the existence of multi-taxa interactions. Further work is required to determine the functionality of these multi-interacting partners and if these interactions represent keystone guilds (Banerjee et al., 2018). The identification of common interaction pairings across several groups indicates that these interactions, and the methods we employed to identify them, are robust and presumably not influenced by physiological state or other environmental conditions.

It is important to highlight that these network maps represent patterns, and not direct interactions. Many of the observed interactions may be due to microbes sharing a similar ecological niche, and a substantial challenge, particularly for highly interconnected taxa, will be to investigate these interactions further. To undertake initial validation steps and to demonstrate that microbial interactions are an important factor influencing the colonization of gut-associated microbiota, we infected six culturable bacterial taxa into *Ae. aegypti* larvae that either possessed or lacked their resident microbiota. *Serratia* and *Cedecea,* which were isolated from *Ae. albopictus,* poorly infected *Ae. aegypti* when it possessed its native microbiome, however, when mosquitoes lacked their native microbiota, these bacteria infected at a higher titer. Even more striking was the effect on prevalence (number of individuals infected), which changed from 38% and 50% for *Serratia* and *Cedecea*-*alb,* respectively, in conventionally reared mosquitoes that possessed their resident microbiota, to 100% infection when infected into axenic larvae. This indicated that the poor infection rates were not related to host or bacterial genetics, but to microbial incompatibility. In our networks, *Serratia* and *Cedecea* have several co-exclusionary relationships with dominant bacterial taxa such as *Asaia, Pseudomonas,* and *Enterobacter,* which may explain these results, however specific examination of these interactions in adult mosquitoes is required. Importantly, not all of the bacterial taxa artificially infected into larvae increased in prevalence and density suggesting these enhanced colonization effects are not simply due to mosquitoes lacking their microbiota, but rather are specific for each taxa, likely due to specific microbial interactions. Similar to our findings, it has been reported that antibiotic treatment prior to bacterial supplementation in a sugar meal can increase the prevalence of infection of gut microbes in female *Anopheles* and *Aedes* mosquitoes (Ramirez et al., 2014), indicating that resident gut bacteria that are susceptible to antibiotics are antagonistic to the supplemented bacterium. It is important to note that our reinfection study, which exploited the gnotobiotic rearing system, examined interactions in larvae, not adults, and that differences in the gut morphology and function between these two life stages may alter microbial interactions (Engel and Moran, 2013). However, our findings combined with the work of Ramirez *et al.* (2014), suggest that microbial interactions between gut-associated bacteria occur within mosquitoes and influence symbiont colonization in aquatic and adult life stages, which likely affects microbiome species richness and evenness. These colonization traits and co-exclusionary associations could offer a possible explanation for the variability seen in the mosquito microbiome between individuals, as bacteria that initially infect the gut may impede colonization by other microbes.

In this study, we compared microbial interaction networks from field and laboratory mosquitoes to examine the influence of the general environment but it is possible other factors may influence network structure. In particular, it would be interesting to determine if microbial networks differ between tissues within mosquitoes, such as the salivary glands, germline and the gut. For example, microbial network analysis from the human microbiome project found strong niche specialization in their networks, whereby different body sites had contrasting microbial networks (Faust et al., 2012). In *Anopheles* mosquitoes, salivary glands have a more diverse microbiome compared to the gut (Sharma et al., 2014), and it is conceivable that elevated species richness would allow for greater network interactions. Furthermore, the germlines of male and female *Anopheles* mosquitoes share some common taxa but there are also quantitative differences (Segata et al., 2016). These differing microbial niches could be exploited to determine the influence of species evenness on microbial networks. Performing network analysis on gut samples of *Ae. albopictus* and *Cx. quinquefasciatus* may also overcome any issue with *Wolbachia* sequestering the majority of the reads, as *Wolbachia* primarily resides within the germline in these mosquito species. Here, we assessed whole mosquitoes to give an initial overall picture of microbial interactions, but analysis of distinct tissues may identify interactions of bacteria that are proximal to one another, and these interactions are more likely to reflect true microbe-microbe interactions, rather than patterns associated with environmental exposure. Within a species we collected mosquitoes from a single site, but future studies examining interaction networks should incorporate diverse sites. Common pairwise interactions identified across sites would indicate robust relationships not influenced by environmental factors.

In summary, we examined the microbiome of three important mosquito vectors, *Ae. aegypti, Ae. albopictus,* and *Cx. quinquefasciatus.* While the overall microbiome structure between host-seeking or ovipositing females was similar, we identified specific bacteria that changed in abundance between mosquito species. Our analysis identified a suite of pairwise interactions used for generating microbial interaction networks, and together with re-infection studies we have demonstrated that microbial interactions affect microbiome composition and abundance of specific bacterial taxa. These findings add to our understanding of microbiome community structure of mosquitoes and factors that influence microbiome acquisition and maintenance in these important disease vectors.

## Acknowledgements

We thank the UTMB insectary core for providing the laboratory mosquitoes. GLH is supported by NIH grants (R21AI124452 and R21AI129507), a University of Texas Rising Star award, the John S. Dunn Foundation Collaborative Research Award, the Robert J. Kleberg, Jr. and Helen C. Kleberg Foundation, and the Centers for Disease Control and Prevention (CDC) (Cooperative Agreement Number U01CK000512). The paper’s contents are solely the responsibility of the authors and do not necessarily represent the official views of the CDC or the Department of Health and Human Services. This work was also supported by a James W. McLaughlin postdoctoral fellowship at the University of Texas Medical Branch to SH, a NIH T32 fellowship (2T32AI007526) to MAS and a Marie Curie Fellowship (within the 7^th^ European Community Framework Programme; project no. 330136) to EAH.

## Data accessibility

The NCBI accession number for the raw sequencing data reported here is PRJNA422599.

## Author Contributions

SH, KK, LA, JAD, EAH, YF, and GLH designed the experiments. GCM, CLF, JAD and MD oversaw the fieldwork. SH and MP completed the experiments. KK, MMR, LA, SH, EAH and GLH undertook analysis. SH, KK, JAD, MD, and GLH wrote and edited the manuscript and all authors agreed to the final version. YF and GLH acquired funding and supervised the work.

## Supplementary figures

**Figure S1**. Map of Houston, Texas, indicating the field collection sites.

**Figure S2**. Shannon entropy rarefied at intervals between 0 and 100,000 reads in each sample from different groups (G, BG, Laboratory) in all three mosquito species.

**Figure S3**. Heat maps indicating bacterial relative abundance for the three mosquito species. OTUs were grouped to genus level or higher ranks (when genus was ambiguous) and the relative abundance indicated by color for each individual (column) is shown. The upper heat map is with *Wolbachia* present while the lower has *Wolbachia* excluded. The dendrogram/clustering of the bacteria is generated based on their relative abundance correlation across samples.

**Figure S4**. Shannon diversity of *Ae. albopictus* and *Cx. quinquefasciatus* with and without *Wolbachia.* For analysis of samples without the endosymbiont, *Wolbachia* reads were computationally excluded from the analysis and then Shannon diversity was recalculated.

**Figure S5**. Relative abundance of *Serratia* and *Aeromonas* from high-throughput sequencing in *Cx. quinquefasciatus, Ae. albopictus* and *Ae. aegypti* mosquitoes reared in the lab. Data were analyzed using a one-way ANOVA using Tukeys method for pairwise comparisons (**** is P<0.0001).

**Figure S6**. Examples of co-occurrence and co-exclusion microbial pairs identified in the interaction networks. Scatterplots of relative abundance profiles displaying statistically significant co-occurrence and co-exclusion patterns in mosquito groups. Points represent the relative abundance values of the pair in each sample.

## Supplemental tables

**Table S1**. Number of samples used in the study and their division into groups by metadata.

**Table S2**. Complete and filtered to 0.1% OTU table with read counts from each individual mosquito (Ae. *aegypti, Ae. albopictus Cx. quinquefasciatus).* Each library was constructed from a single female mosquito.

**Table S3**. List of the bacterial taxa present or absent within each species and group which was used to create Venn diagrams (Figure 3A).

**Table S4**. OTU tables (relative abundance profiles) used in CoNet analysis. Relative abundance of each OTU in a sample was calculated by dividing number of reads by total number of reads of the sample and OTUs with read counts below 0.1% across all mosquito samples were excluded from analysis. OTU relative abundance profiles were summed based on lowest common taxonomy, down to genus level. When building networks for each mosquito group (Ae. *aegypti* BG, G or Lab, *Ae. albopictus* BG, G or Lab, or *Cx. quinquefasciatus* BG, G or Lab), abundance profiles present (non-zero abundance) in less than 5 samples were excluded from analysis.

